# Did position-effect guide the evolutionary dynamics of developmental gene expression?

**DOI:** 10.1101/127563

**Authors:** Meenakshi Bagadia, Keerthivasan Raanin Chandradoss, Yachna Jain, Harpreet Singh, Mohan Lal, Kuljeet Singh Sandhu

**Author notes:** Equal contribution (Meenakshi Bagadia, Keerthivasan Raanin Chandradoss). To whom correspondence should be addressed: Kuljeet Singh Sandhu, Assistant Professor, Department of Biological Sciences, Indian Institute of Science Education and Research (IISER) - Mohali, E. mail.

## Abstract

Conserved noncoding elements (CNEs) have significant regulatory influence on their neighbouring genes. Loss of synteny to CNEs through genomic rearrangements can, therefore, impact the transcriptional states of the cognate genes. Yet, the evolutionary implications of such chromosomal position effects have not been studied. Through genome-wide analysis of CNEs and the cognate genes of representative species from 5 different mammalian orders, we observed significant loss of synteny to CNEs in rat lineage. The CNEs and genes losing synteny had significant association with the fetal, but not the post-natal, brain development as assessed through ontology terms, developmental gene expression, chromatin marks and genetic mutations. The loss of synteny correlated with the independent evolutionary loss of fetus-specific upregulation of genes in rat brain. DNA-breakpoints implicated in brain abnormalities of germ-line origin had significant representation between CNE and the gene that exhibited loss of synteny, signifying the underlying developmental tolerance of genomic rearrangements that had allowed the evolutionary splits of CNEs and the cognate genes in rodent lineage. These observations highlighted the non-trivial impact of chromosomal position-effect in shaping the evolutionary dynamics of mammalian brain development and might explain loss of brain traits, like cerebral folding of cortex, in rodent lineage.

**Author Summary:** Expression of genes is regulated by proximally located non-coding regulatory elements. Loss of linear proximity between gene and its regulatory element thus can alter the expression of gene. Such a phenomenon can be tested at whole genome scale using evolutionary methods. We compared the positions of genes and regulatory elements in 5 different mammals and identified the significant loss of proximities between gene and their regulatory elements in rat during evolution. Brain development related function was selectively enriched among the genes and regulatory elements that had lost the proximity in rat. The observed separation of genes and their regulatory elements was strongly associated with the evolutionary loss of developmental gene expression pattern in rat brain, which coincided with the loss of brain traits in rodents. The study highlighted the importance of relative chromosomal positioning of genes and their gene regulatory elements in the evolution of phenotypes.

## Introduction

Around 4-8% of the human genome is evolutionary constrained, of which coding elements contribute only about 1.5%, while rest is non-coding (1-3). Massive data produced by ENCODE and Epigenome Roadmap projects have confirmed that majority of the evolutionary constrained non-coding DNA serve as protein binding sites(4, 5). These conserved noncoding elements (CNEs) are interwoven with the protein coding genes in a complex manner. Ample evidence converges to non-trivial regulatory impact of CNEs on proximal gene. Deletion of a non-coding region between sclerostin (SOST) gene, a negative regulator of bone formation, and MEOX1 impacts the expression of SOST and is strongly associated with Van Buchmen disease characterized by progressive overgrowth of bones (6). Similarly, deletion of a 10kb non-coding region downstream to stature homeobox (SHOX) gene is associated with Leri Weill dyschondrosteosis syndrome, a skeletal dysplasias condition (7). Mutations in CNEs downstream to PAX6 gene prevent its expression and are associated with Aniridia, a congenital eye malformation. Genetic errors in locus control region (LCR) at alpha and beta globin loci strongly associate with alpha/beta-thalassemia (8, 9). Maternal deletion of Igf2/H19 ICR disrupts the Igf2 imprinting leading to bi-allelic expression of Igf2, which is strongly associated with Beckwith Weidman syndrome (10). Loss of a CNE proximal to androgen receptor is strongly associated with evolutionary loss of penile spines and sensory vibrissae in human (11).

Around 200,000 human-anchored Conserved Non-coding Elements (CNEs) have been identified in mammals, which are likely to exhibit gene regulatory potential, as measured through enhancer-associated chromatin marks (12-14). Most CNEs position around developmental genes (14-16). However, establishing causal relationship between CNE and the phenotype remains a daunting task. Though genome wide association studies (GWAS) have uncovered a whole repertoire of non-coding variants with phenotypic associations (17), it is difficult to identify the causal variants. More recently, pooled CRISPR-Cas technique has been implemented to alter the non-coding elements to assess their function more precisely (18). These methods are difficult to be scaled up for high throughput genotype-phenotype associations. With the availability of whole genome sequences of multiple species, evolutionary methods are instrumental in deciphering genotype-phenotype associations. Through comprehensive multi-species comparison, it has been inferred that most CNEs are syntenic to the nearest gene in linear proximity and are likely to regulate the same (14, 19). Attempts have been made to link evolutionary loss and sequence divergence of CNEs to lineage specific traits, like auditory system in echo-locating mammals and adaptively morphed pectoral flippers in marine mammals (20, 21). In this study, we asked the question whether the lineage-specific evolutionary alterations in relative chromosomal positions of CNEs are associated with lineage-specific changes in gene expression. Through analysis of chromosomal positions of orthologous CNEs and genes from 5 different mammals, we observed that a significant number of genes had lost synteny to their adjacent CNEs independently in rat lineage. This loss of synteny was significantly associated with the down-regulation of genes involved in neurogenesis and neuronal migration during fetal brain development, which coincided with the evolutionary loss of several brain traits independently in rat lineage. The study suggested significant contribution of chromosomal position-effect in the evolutionary divergence of developmental gene expression trajectories in mammals.

## Results

### Loss of synteny between CNE and the proximal gene

Using chromosomal position data of CNEs and genes from representative primate (human), rodent (rat), carnivore (dog), perrisodactyl (horse), and artiodactyls (cow), we obtained 51434 ‘syntenic’ CNE-gene pairs (4241 genes), wherein the CNE and the nearest gene-TSS were <1 Mb distance apart in all 5 species. There were 3579 ‘non-syntenic’ CNE-gene pairs (334 genes), wherein the CNE and the gene-TSS were on different chromosomes or were >2Mb apart independently in one of the species (Figure 1A, Figure S1, Methods). The rationale of 1Mb distance cutoff for the synteny was based on the observation that the distribution of all CNE-gene distances saturated when approached 1Mb range (Figure 1B). Similar approach has been used earlier to infer enhancer-promoter linkage based on evolutionary syntney between the two in 1Mb range(19). The distance cut-off of 2Mb for loss of synteny made sure that the minimal expansion in CNE-gene distance would at-least be 2 fold. To test if the CNEs in syntenic and non-syntenic sets were comparable, we assessed their lengths and degree of conservation in mammalian genomes. Figure 1C showed insignificant differences in the degree of sequence conservation of syntenic and non-syntenic CNEs, suggesting that the sequence of non-syntenic CNEs had not diverged among mammals as compared to that of syntenic CNEs. The length distribution of syntenic and non-syntenic CNEs showed only marginal difference towards slightly longer CNEs in non-sytenic set (Figure 1D). However, the syntenic and non-syntenic CNEs were located in the genomic domains of distinct sequence properties. We analysed the enrichment of SINE, LINE and LTR retrotransposons, which covers upto 50% of mammalian genome, around syntenic and non-syntenic CNEs. Syntenic CNEs were enriched in the region of open chromatin, as signified through greater enrichment SINE content, and might have more wide-spread role across different cell-lineages as compared with the non-syntenic CNEs (Figure 1E). In contrast, the non-syntneic CNEs were located in the domains enriched with long terminal repeats (LTR), marking their susceptibility to genomic rearrangements through mechanisms like non-allelic homologous recombination (NAHR) (22, 23) (Figure 1E). LINE elements, in general, did not exhibit significant difference in the two sets. These observations were largely consistent across species, marking an ancestral property (Figure 1E), except in the rat wherein the LINE elements were enriched around non-syntenic CNEs. This exception can be explained by the fact that the rodents retained least of the ancestral retrotransposons as compared to most other mammals and had accumulated newer elements(24).

**Figure 1.**
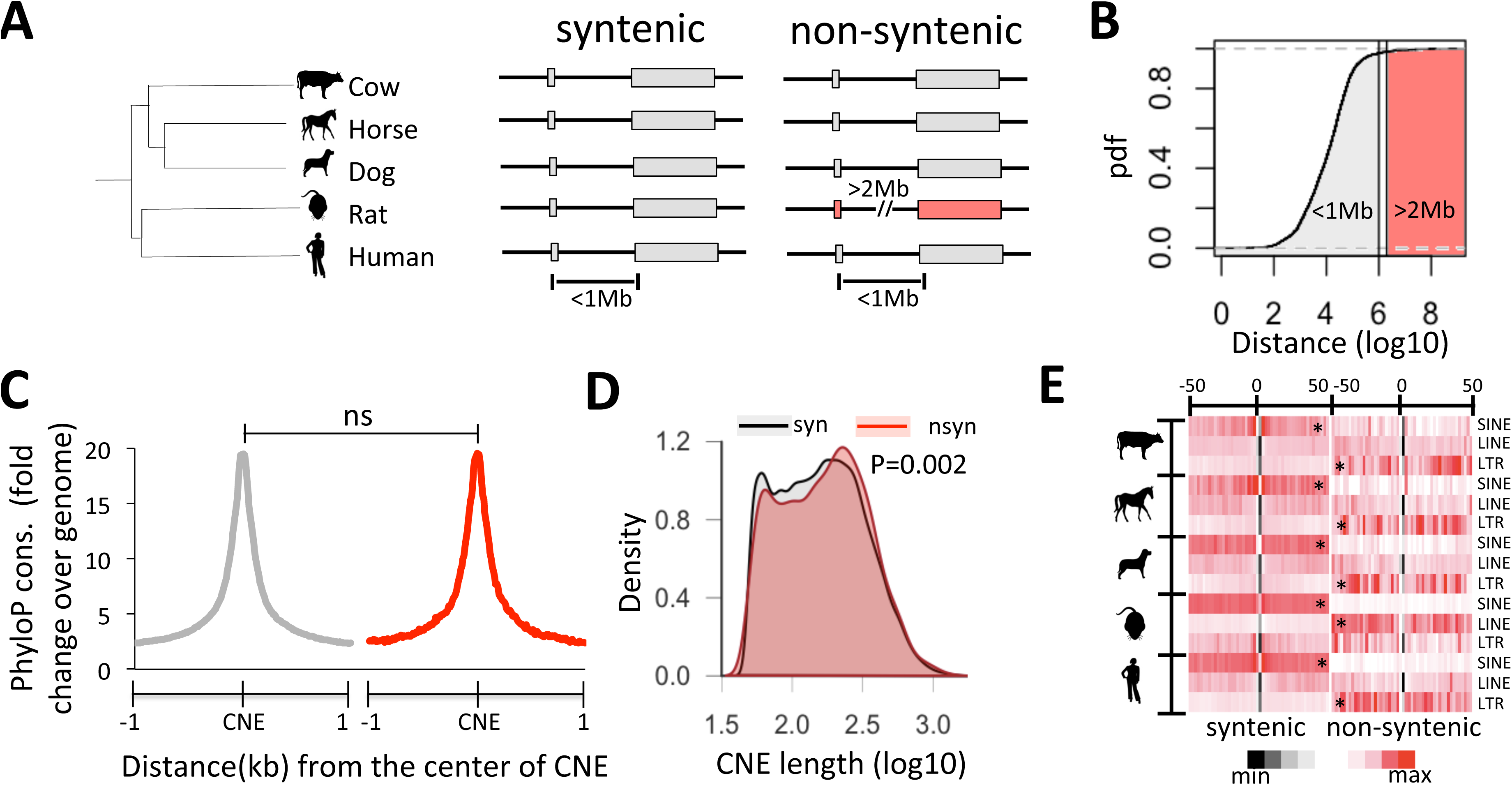
Synteny and lack thereof between CNE and the nearest gene. (a) Illustration of the strategy to infer the synteny and lack thereof between CNE and the neighboring gene across 5 representative mammalian orders. CNE-gene pairs were classified as syntenic if remained proximal (<1Mb) in all the 5 species and as non-syntenic if departed by > 2mb or on different chromosomes in on of the species while maintaining synteny in other 4 species. (b) Cumulative distribution of all CNE-gene distances in the human genome. Most CNE-gene pairs were <1Mb distance and, therefore, cut-off of 1Mb was applied for CNE-gene synteny. (c) Sequence conservation, as measured through mammalian PhyloP scores, and (d) length distribution of CNEs in syntenic and non-syntenic sets. (e) Enrichment of retrotransposons +/− 50Kb around syntenic and non-syntenic CNEs. Asterisk indicate significant p-values (<0.05) calculated using Mann Whitney U test of enrichment values +/− 10kb around CNEs.

Relatively large number of syntenic CNE-gene pairs (93.5%) confirmed the widespread conservation of linear proximity between CNE and its adjacent gene (14). Among the total 3579 non-syntenic instances, 2711 (75%) were associated with the rat genome alone, coherent with the significantly greater number of structural variations in rodent clade(25) (Figure 2A-B). Positive scaling between number of non-syntenic instances and the break-point distances of species from the common ancestor signified that CNE-gene synteny was an ancestral trait (Figure 2B). Due to significant loss of synteny in rat lineage as compared to others, we focussed on rat instances in this study. By ‘Loss of synteny’ or ‘non-syntenic’ set, we referred to loss of CNE-gene synteny in rat from figure 2C onwards. To directly assess the proportion of non-syntenic CNE-gene pairs associated with structural variations, we analysed the rodent-specific evolutionary break-points (Methods). We observed that 930 (34%) of all non-syntenic instances in rat had at-least one rodent-specific break-point in between the gene-TSS and the CNE as compared to 319 (11%) on an average for the random null prepared through distance-controlled bootstrap sampling of syntenic CNE-gene pairs (Figure 2C, Methods). This suggested that the loss of CNE-gene synteny in rat could largely be explained through rodent-specific genomic rearrangements (Figure 2C). We further argue that the sequence alignment based annotations of evolutionary breakpoints might not represent the entire repertoire of genomic rearrangements and, therefore, analysed the neighbouring genes on either side of non-syntenic CNEs to map the various rearrangement scenario through which CNE-gene synteny was lost. We found that that the translocation like scenario, as marked by (i) in Figure 2D, largely explained the inter-chromosomal (*trans*) splits of CNE and the adjacent gene. The scenario, which reflected the mapping artefacts, like in panel (iii), was under-represented (5%, Figure 2D). Analysis of intra-chromosomal (*cis*) splits suggested inversion-like events separating the CNE-gene pairs. Scenario (iv) and (v) showed events where region adjacent to CNE (on left side in scenario-iv and right side in scenario-v) had undergone local rearrangements, of which 30% and 90% events respectively were confirmed as inversion events by analysing the change in relative strand orientation of neighboring genes. We illustrated examples of *trans* and *cis* splits of CNE and the genes in Figure 2E. Gene POU3F2 on human chromosome 6 was syntenic to a CNE, which was 45Kb upstream. The orthologous CNE and the gene in rat were on chromosome 8 and 5 respectively marking the *trans* split of CNE and the gene through translocation (Figure 2E). Another CNE was 18kb upstream to gene ADAM23 on human chromosome 2. The rat orthologues were separated by 2.4Mb on chromosome 9 through an inversion (Figure 2E).

**Figure 2.**
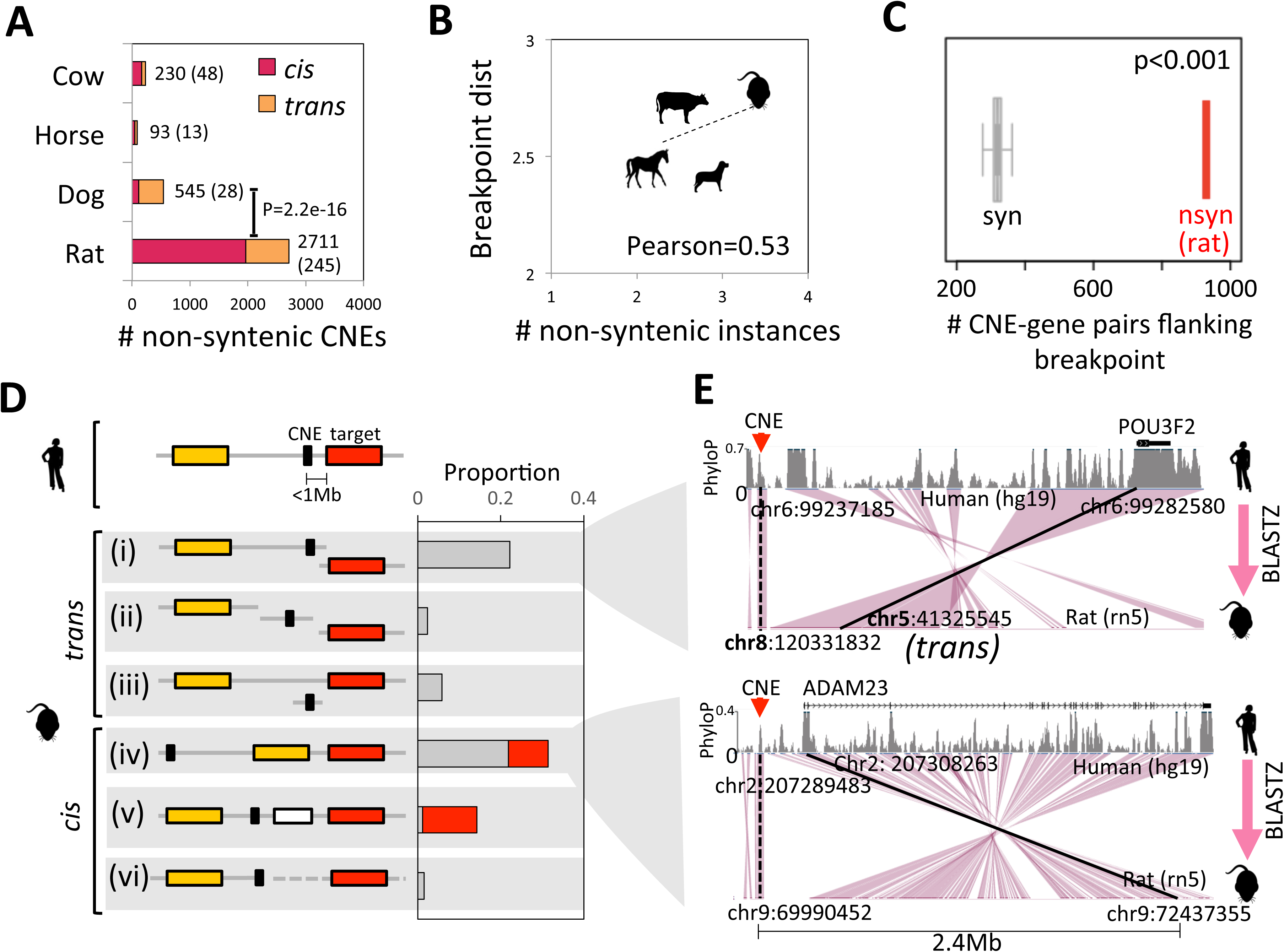
Genomic rearrangements underlying the loss of synteny. (a) Number of non-syntenic instances and genes (in parentheses) in different mammals. P-value for the non-syntenic instances in rat and the next highest value (in dog) was calculated using Fisher’s exact test. (b) Scaling between number of non-syntenic cases and the evolutionary break-point distance from the common mammalian ancestor. (c) Number of non-syntenic CNE-gene pairs having atleast one rodent-specific break-point inbetween CNE and gene-TSS, overlaid onto null distribution prepared from syntenic set. (d) Distinct trans and cis chromosomal rearrangements as inferred from the analysis of genes flanking the non-syntenic CNEs. Shown are the neighboring genes around CNE. Red color represents the target gene and orange color represents the nearest gene on the other side of the CNE. Red color in the barplot marks the proportion for which inversion could be confirmed through analysis of gene orientations. Shown are the two examples illustrating loss of CNE-gene synteny in rat. In first example, a CNE was located 45kb upstream to the gene POU3F2 in human, but were split on different chromosomes in rat. Second example shows that an inversion event had distanced the CNE and the proximal ADAM23 gene upto 2.4Mb in rat genome.

We concluded that the rodent-specific genomic rearrangements largely explained the loss of CNE-gene synteny in rat.

### Genes that had lost synteny to CNEs in rat were associated with the fetal brain development

Significant differences in the genomic attributes around syntenic and non-syntenic CNEs hinted at their distinct functional roles. To assess their functions, syntenic and non-syntenic gene-lists were subjected to Gene Ontology (GO) and Mammalian Phenotype Ontology (MPO) analyses. The analysis of GO terms revealed enrichment of general as well as various tissue-specific development related terms in the syntenic set, while non-syntenic set was specifically enriched with nervous system development related terms (Figure 3A). In MPO analysis, syntenic set exhibited of enrichment of neonatal lethality and skeletal phenotypes, while non-syntenic set was associated with brain morphology related phenotypes (Figure S2A). Species, other than rat, did not exhibit enrichment of any particular functional term, owing to smaller sample size.

**Figure 3.**
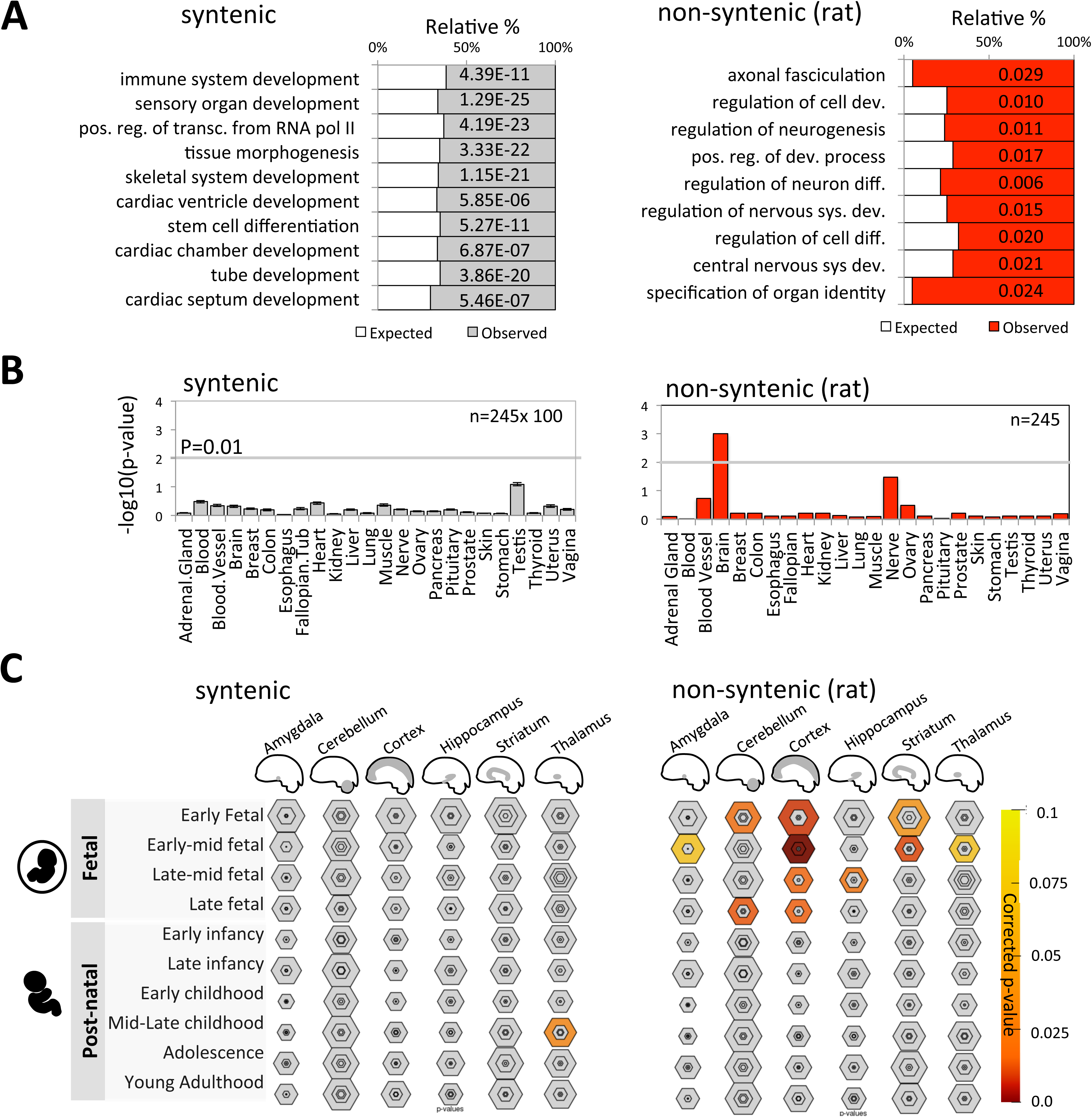
Functional characterization of genes in syntenic and non-syntenic sets. (a) Enrichment of Gene Ontology (GO) terms among genes in syntenic and non-syntenic sets. P-values shown were corrected using Benjamini-Hochberg method. (b) Tissue-specific expression analysis of genes in syntenic and non-syntenic set. Relative significance was plotted as negative of log10 transformed corrected p-values of Fisher’s exact test for the overlap with the tissue-specific genes at stringency score (pSI) < 0.05. Horizontal grey colored dashed line represents the p-value of 0.01. For syntenic set, mean values and standard errors of significance for 100 random samples of 245 genes (the size of non-syntenic set) from syntenic sets were plotted. (c) Expression specificity of genes in syntenic and non-syntenic sets across brain regions and across developmental stages. For syntenic set, the sample that exhibited maximum significance for brain specificity in panel-b was taken. Size of the nested hexagons represents the proportion of all genes specifically expressed in particular tissue at particular developmental stage. Hexagons are nested inwards based on relative stringency of tissue specificity scores (pSI=0.05, 0.01, 0.001 & 0.0001 respectively). Color gradient represents the magnitude of corrected p-values of Fisher’s exact test.

We further followed the above observations through tissue-specific gene expression analysis in human. The syntenic set had widespread representation of genes expressed in different cell-lineages and, therefore, did not exhibit significant tissue-specificity, while genes in non-syntenic set were specifically expressed in brain (Figures 3B). The brain-specific expression of genes in non-syntenic set was also confirmed through enrichment analysis of anatomical terms from *bgee* database (Figure S2B). Within brain, non-syntenic set was enriched with the genes specifically expressed in cerebral cortex during fetal, but not post-natal development (Figure 3C). In contrast, the genes in syntenic set did not exhibit any specificity for brain tissues and developmental stages (Figures 3C). These observations highlighted fetal brain-specific roles of genes in the non-syntenic set.

### Non-syntenic CNEs function as fetal brain-specific enhancers

To test whether the differences between syntenic and non-syntenic sets observed through functional analysis of genes, were coherent with the associated CNEs, we tested the regulatory potential of CNEs by analyzing their epigenomic properties across tissues. Through analysis of enhancer associated chromatin state annotations from Epigenome Roadmap, ENCODE and Fantom consortia (Methods), we observed that 74% of syntenic and 61% of non-syntenic CNEs overlapped with the enhancer-associated regulatory sites in at-least one of the tissues or cell-types, marking the enhancer potential of CNEs. Relatively less representation of enhancers in the non-syntenic set might relate to their tissue or developmental stage specific functions, a hypothesis that we further reconciled through detailed analysis of histone modification associated with enhancers, namely Histone-3-Lysine-4-mono-methylation (H3K4me1). We chose this mark because of its strong association with the enhancer potential and the availability of genome-wide datasets for all the cell-lineages we were interested in. We observed that: 1) the CNEs in syntenic set exhibited consistent H3K4me1 enrichment across several fetal and adult tissues like thymus (endodermal), muscle (mesodermal), heart (mesodermal), intestine(endodermal) and brain (ectodermal) (Figure 4A, Figure S3A-B); 2) H3K4me1 enrichment over CNEs in non-syntenic set was specifically higher (comparable to that of syntenic CNEs) in fetal, but not in adult brain (Figure 4A). We further observed the significant enrichment of binding sites of ectoderm-specific transcription factors, which were specifically upregulated in fetal brain, in non-syntenic CNEs as compared to syntenic CNEs (Figure S4, Methods). These observations were largely coherent with our proposal that non-syntenic CNE-gene pairs were associated with fetal brain development.

**Figure 4.**
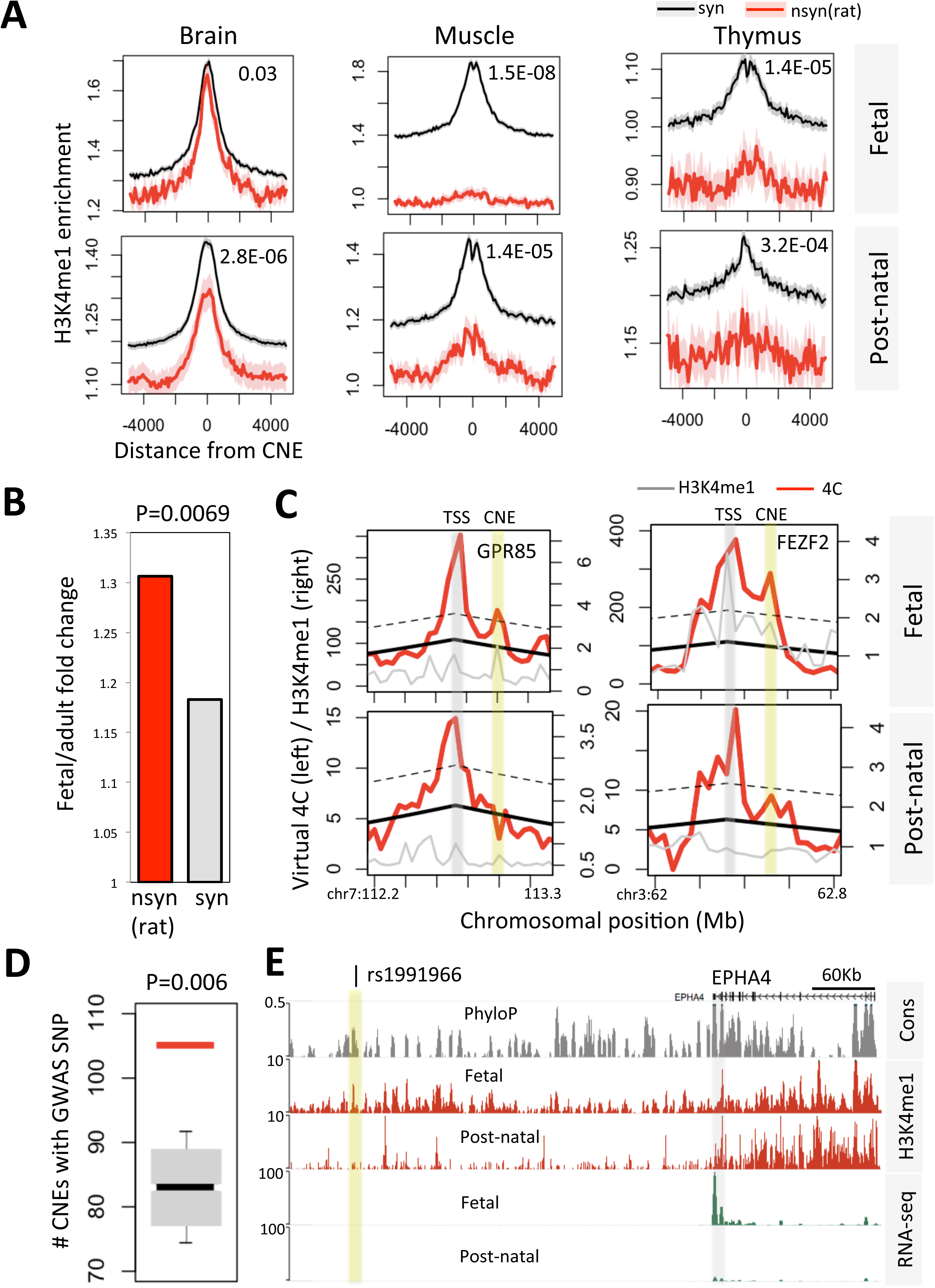
Enhancer properties of syntenic and non-syntenic CNEs. (a) H3K4me1 ChIP enrichment (over input) on and around CNEs in syntenic and non-syntenic sets in fetal and post-natal tissues. P-values for the difference between syntenic and non-syntenic CNEs were calculated using Mann Whitney U test of H3K4me1 enrichment values in 1kb spanning windows on either side of the CNEs. (b) Virtual 4C analysis of CNE-gene interactions in fetal and adult human brains. The barplot shows fetal-to-adult ratio of proportion of CNE-gene pairs exhibiting significant physical chromatin interactions. (c) The examples of GPR85 and FEZF2 genes are shown for illustration. Vertical grey and yellow bars represent the TSS and CNE positions respectively. Red and grey curves show virtual 4C and h3K4me1 signals respectively. Black smooth line represents the loess fit of the 4C signal as function of genomic distance from the reference point. The dotted line represents 3σ distance from the loess regression line. (d) Proportion of non-syntenic CNEs having at-least one brain associated SNP superimposed onto the null distribution obtained from syntenic CNEs. P-value was calculated using boot-strap method by randomly sampling 2711 CNEs (size of non-syntenic set) from syntenic set 1000 times. (e) An example of EPHA4 gene and its proximal CNE having a Schizophrenia associated GWAS SNP is shown. The tracks for PhyloP conservation score, H3K4me1 and RNA-seq data of fetal and post-natal brain aligned accordingly. The orthologous CNE and gene were 7.2 Mb apart on chromosome 9 in rat.

To assess the physical enhancer-promoter association, we generated virtual 4C data by processing available HiC datasets of fetal and adult human brains (Figure S3C). Figure 4B showed the significant fetal-to-adult ratio of the proportion of non-syntenic CNE-gene pairs showing significant physical interactions as compared to that of syntenic CNE-gene pairs. We illustrated the physical interactions between CNE and the gene through examples (Figure 4C, Figure S3D). Transcription start sites of GPR85 and FEZF2 genes, both associated with neurological phenotypes(26-28), showed significant interaction frequency to their cognate CNEs in human fetal brain, but not in adult brain. The H3K4me1 signals at TSSs and CNEs were also significant in fetal brain as compared to adult. Epigenomic analyses thus suggested that the majority of the non-syntenic CNEs exhibited enhancer associated hallmarks in fetal brain.

By mapping the trait/disease associated SNPs from Genome Wide Associated Studies (GWAS) and the nearby SNPs (proxy) in the linkage disequilibrium based on 1000 genome data, we observed that 105 of the non-syntenic CNEs were having at-least one brain related SNP (Figure 4D). This representation was statistically significant when compared with that of syntenic set (Figure 4D). These observations represented genetic evidence of brain-specific roles of non-syntenic CNEs. We highlighted the example of EPHA4 gene, which is required for radial neuron migration and is involved in the pathways leading to lissencephaly and schizophrenia in human(29, 30). Upstream CNE to this gene had a schizophrenia associated SNP. Fetal brain specificity of CNE and gene expression was illustrated using H3K4me1 and RNA-seq tracks of fetal and post-natal human brains (Figure 4E).

Therefore, our observations through enhancer datasets, epigenomic marks, differential motif enrichment analysis and brain associated SNPs concomitantly established that the non-syntenic CNEs were specific to fetal brain development in human.

### Developmental tolerance loss of synteny events

While we have shown that the genes and the CNEs that had lost synteny in rat were associated with fetal brain development, whether or not CNE-gene proximity was causally linked with the brain-specific expression of the cognate gene remained to be addressed. Towards this, we assessed the representation of germ-line breakpoints associated with the congenital disorders exhibiting brain abnormalities and the somatic cancer breakpoints, between CNE and gene-TSS in syntenic and non-syntenic sets. Since the observed germ-line break-points are the ones that had survived through germ-line and the embryonic development, their presence and absence between CNE and the adjacent gene signifies developmental tolerance and intolerance respectively of loss of CNE-gene synteny. On the contrary, the cancer breakpoints of somatic origin do not undergo such selection and hence do not indicate the developmental tolerance or lack thereof. Figure 5A showed relative proportion of non-syntenic CNE-gene pairs having at-least one DNA breakpoint between gene-TSS and the CNE superimposed onto random null prepared from syntenic CNE-gene pairs of similar distance distribution as that of non-syntenic. We observed significantly greater representation of germ-line breakpoints in non-syntenic set as compared to syntenic set, while representation of somatic breakpoints showed insignificant difference (Figure 5A). We interpreted that DNA breakpoints between non-syntenic CNE and the genes were developmentally tolerable and genomic rearrangements thereof in the ancestral genome might have served as an evolutionary substrate for position effect. We further showed a few examples of germ-line chromosomal rearrangements that had split the CNE and the adjacent gene in congenital disorders with brain abnormalities (Figure 5B). Example (i) in Figure 5B showed a chromothripsis event wherein an inversion had split the CNE-gene pair. The involved gene BCL11A regulates cortical neuron migration and mutations therein associate with microcephaly and intellectual disability in human(31). BCL11A gene also exhibited 3.6 fold loss of expression in peripheral blood of the patient having genomic rearrangement as compared with the normal mother of the patient. Example (ii) showed a translocation event splitting a CNE and CCDC68 gene. Genetic mutations in CCDC68 are associated with schizophrenia, bipolar disorder and autism(32). In example (iii), an inversion had split the CNE from DNAJB6 gene, which has role in neuritogenesis and neuroprotective functions(33).

**Figure 5.**
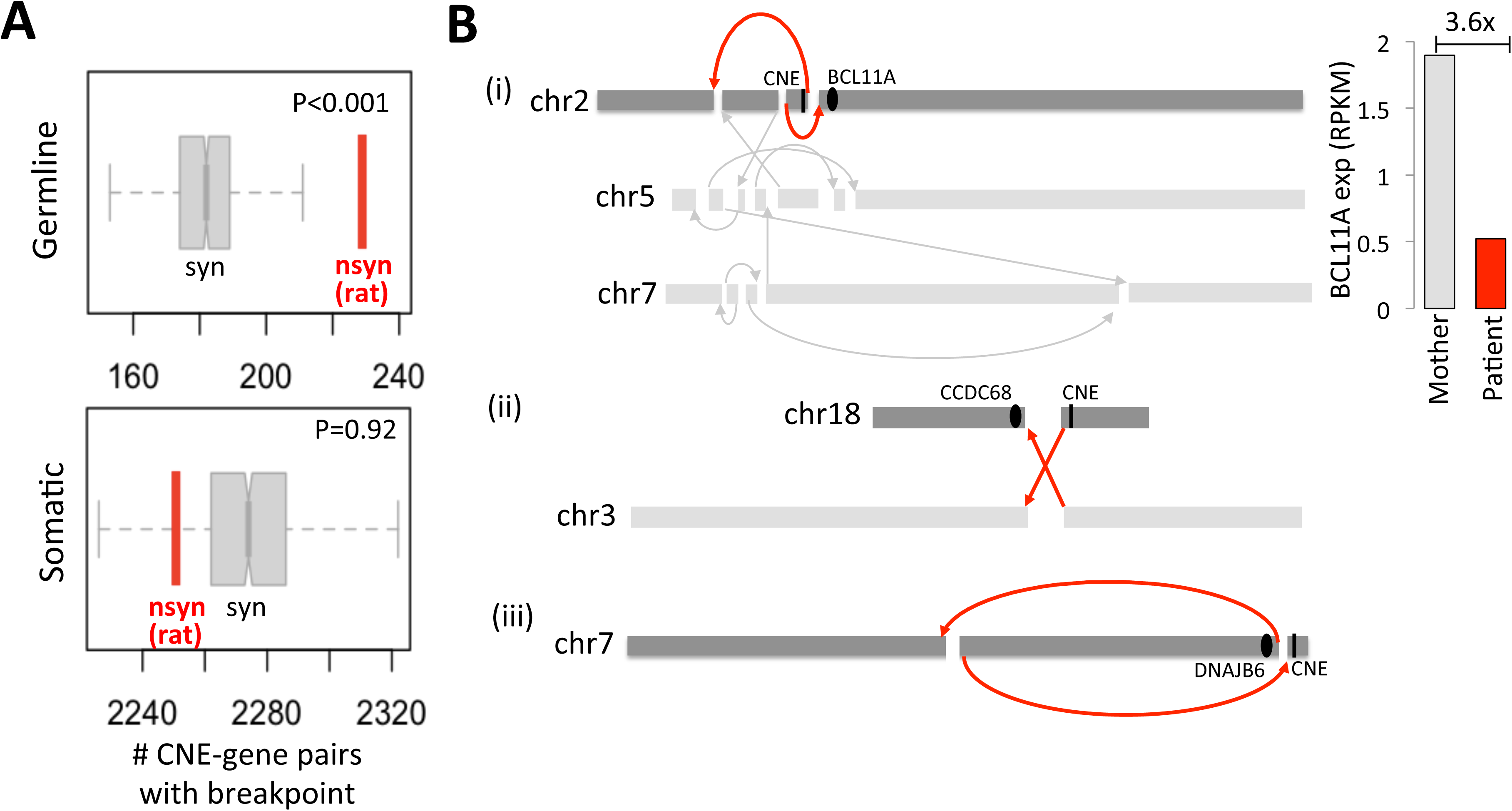
Tolerance and intolerance of CNE-gene split. (a) Number of non-syntenic CNE-gene pairs flanking at-least one germ-line breakpoint associated with the congenital disorders having brain abnormalities (upper panel) and somatic cancer breakpoint (lower panel), superimposed onto null distributions obtained from the syntenic set of same CNE-gene distance distribution as that of non-syntenic set. P-values were calculating using boot-strap method with 1000 random samplings. (b) Examples illustrating chromothripsis, translocation and inversion events breaking CNE-gene synteny in congenital disorders with brain abnormalities. Mutations in BCL11A (i), CCDC68 (ii) and DNAJB6 (iii) genes are associated with abnormal brain phenotypes as elaborated in the results section. The right most panel shows difference in expression level of BCL11A gene in the patient having genomic rearrangement and the normal mother.

### Loss of synteny to CNEs coincided with the loss of fetus-specific upregulation of genes in rat brain

An important question was whether or not the evolutionary loss of CNE-gene synteny in rat was associated with the loss of expression. To assess the functional fate of associated genes, we compared their time-course gene expression trajectories for developing cerebral cortex of human, rat and sheep (as an out-group). Sheep was inducted in the analysis due to the availability of gene expression datasets for pre- and post-natal tissues. We found that 99.4% of CNE-gene pairs that had lost synteny in rat were syntenic in sheep too, confirming the independent loss of synteny in rat lineage. We observed relative loss of fetus-specific gene expression in rat brain as compared to that of human and sheep, suggesting that the loss of synteny correlates with the loss of fetus-specific gene expression in developing rat brain (Figure 6). Enrichment of neurogenesis related genes and down-regulation thereof in fetal brain of rat has implications in understanding loss of brain traits in rat lineage, as discussed in detail in the discussion section.

**Figure 6.**
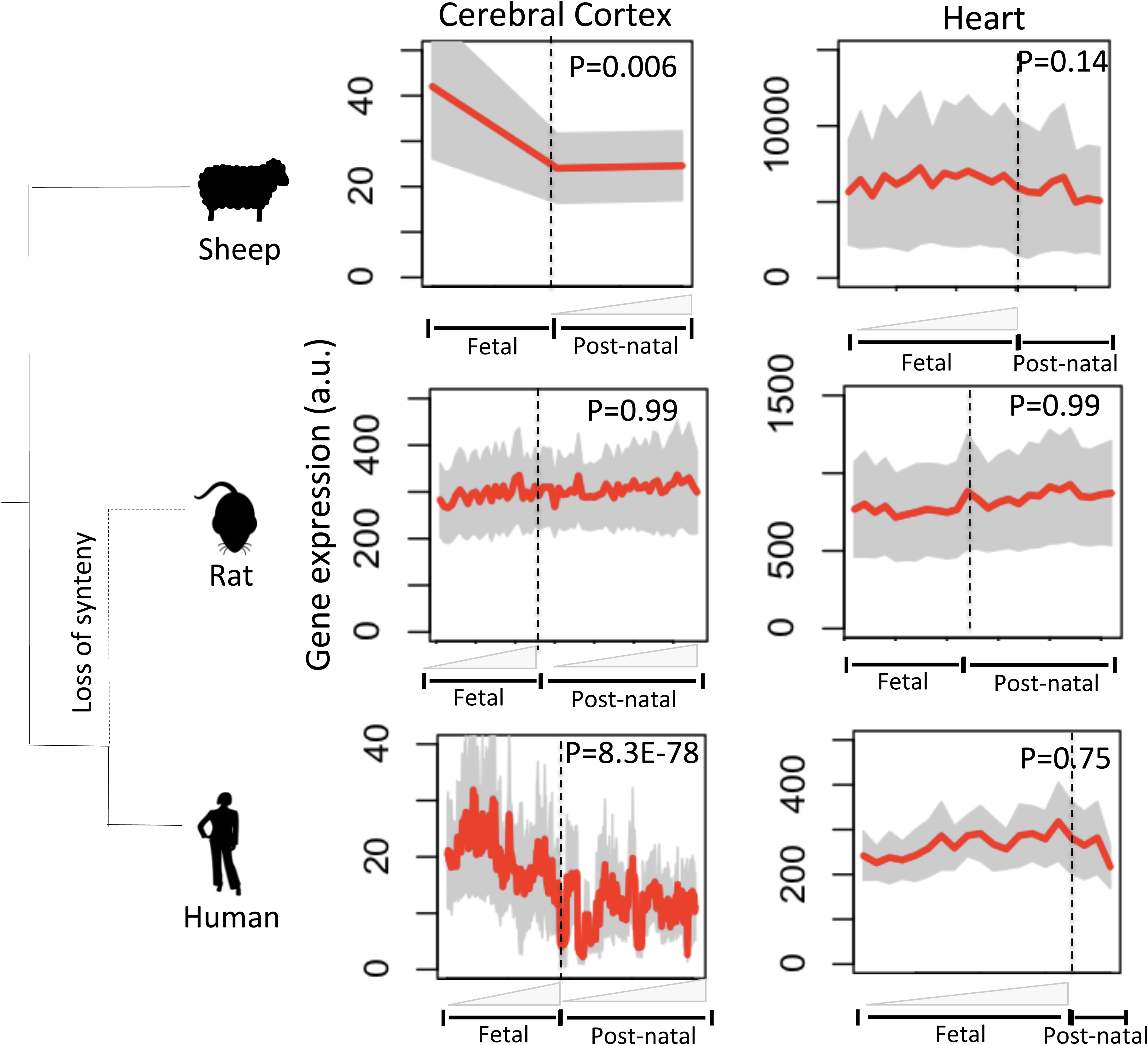
Evolutionary dynamics of developmental gene expression associated with the loss of CNE-gene synteny. Red curves in the plots represent the mean expression of genes in the non-syntenic set and grey area represents 95% confidence interval. Grey colored elongated triangles below the x-axis represent the fetal and post-natal time course. Lack of triangles at some places denotes unavailability of multiple time points in the data. Left panel represents cerebral cortex and right panel is for heart datasets (control).

Taken together, our analysis suggested a strong association between evolutionary dynamics of chromosomal positions of gene regulatory elements and the gain or loss of gene expression, aligning to the notion of ‘position effect’. Tissue and developmental stage specific impact of position effect highlighted the possibility of its significant role in altering developmental dynamics towards evolutionary gain or loss of lineage-specific traits.

## Discussion

It is not always the change in number and the sequence of protein coding regions in the genome that leads to the phenotype alternation in evolution, the dynamics of gene expression is equally relevant in the context. One way the gene expression is altered is through position-effect, i.e., relative chromosomal position of the gene in the genome can alter its expression through regulatory elements and chromatin states in the neighbourhood. Position effect was first discovered through the observation that the chromosomal arrangement of duplicated copies of *bar* gene in *bar*-mutant flies had influence on its expression and consequently causes the relative decrease in number of eye facets (34, 35). Similarly, *white* gene when localized near heterochromatin gives mottled eye phenotype with red and white patches in drosophila eye (36, 37). Despite its significance, the role of position effect in evolution of traits has not been investigated thoroughly. Through comparative genomic analysis, we showed that the CNE-gene pairs that were syntenic in most mammals, but lost the close linear proximity independently in rat were associated with the alteration in the transcriptional program during fetal brain development, presenting evidence how the position-effect might have impacted the evolution of lineage-specific phenotypes by modulating the developmental trajectories in early stages.

Enhancers can function at distance longer than several Mbs and spatial synteny has been observed among genomic regions that had been rearranged in the evolution(38, 39). How might then the loss of linear proximity to CNEs downregulate the expression of genes? Position effect significantly alters the expression noise of the genes(40). Evidence also suggests that long-range or trans enhancer promoter interactions occur at the cost of increased expression noise(41-43). As a result, the overall expression level in a tissue is expected to decline due to increased stochastic fluctuations in gene expression across cells. Therefore, we hypothesized that the loss of linear proximity between CNE and the gene would have compromised with the expression level of the gene by allowing stochastic variations in enhancer-promoter interactions.

Enrichment of brain development related genes in the non-syntenic set might relate to developmental plasticity of brain as compared to other tissues. Genomic alterations at the loci important for the development of basic body plan and functioning would be embryonic lethal, which largely explained the significant representation of skeletal/heart development and neonatal lethality related genes in the syntenic set (Figure 3A, S2A). Brain, despite having neurodevelopment plasticity, exhibits least genome-wide expression divergence across mammalian species(44, 45), but within the space of small non-syntenic gene-set the expression divergence was observed. This suggested that the least expression divergence observed for brain were due to cellular functions that needed to be precisely regulated to maintain delicately shaped brain tissues of all the mammals in general, while the ones that exhibited divergence would implicate in developmental functions specific to fetal brain. Our data showed that one of the ways, such expression divergence was modulated in the evolution was through alteration of genomic proximity between CNEs and the neighbouring genes. Fetus brain-specific downregulation of neurogenesis related genes that had lost synteny to CNEs in rat aligned to the hypothesis that observed genomic alterations might link to brain traits that were lost in rodent lineage. We showed evidence that among the species taken in this analyses, rat exhibited most number of independently modified brain traits, including the ones directly associated with neurogenesis, like absence of cerebral folding of cortex, absence of claustrum separation from cortex, absence of lateral geniculate nucleus magnocellular layer etc. (Figure S5). Of these, loss of cerebral folding of cortex, i.e., lissencephalic or smooth brain phenotype is the largest visible alteration in the rodent brains. Folded or gyrencephalic brain, in general, is considered as adaptation for the mammals with greater encephalization quotient, intelligence and complex behavioural traits(46, 47). It can, therefore, be contended that the CNE-gene proximity and the associated fetal brain-specific expression was not lost in rat, but were rather gained in other mammals that had bigger and gyrencephalic brains. However, we argue that significant non-uniformity in cerebral cortex has been observed across several different mammalian species(48) and the assumption that the common ancestor of placental mammals had a smaller and simpler brain has been challenged recently(49, 50). Evidence converges to gyrencephalic brain of eutherian ancestor and the subsequent loss of cortical gyration in some lineages including rodents has been supported(51). The enrichment of genes associated with brain morphology phenotypes (Figure S2), ECM-receptor interactions & actin cytoskeleton regulation (ACTN, ITGA1, ITGA11, ASAP1, LAMB4, CD36 etc), and the ones implicated in human cortical malformations (MYCN, NRXN1, RASA1, DDX11, FEZF2, EFNA5, GLI3 etc.) (52) in the non-syntenic set further supported the loss of synteny in rat rather than gain of synteny in other mammals (Figure S6). We also emphasized that the loss of synteny in rat was inferred by filtering the CNE-gene pairs which were syntenic in all other species, hence were evolutionarily constrained, except in rat. Assessing gain of synteny was difficult because a CNE-gene pair that was non-syntenic in all species except one cannot be considered as evolutionarily constrained CNE-gene pair. We suspected that gain of synteny inferred in this flawed manner would not have shown any functional association. This indeed was observed through an independent analysis (Figure S6).

It remains arguable whether or not the alterations in brain traits in rodent lineage represented the adaptive selection or was a product of neutral drift. Some studies have suggested that smaller and lissencephalic brain was adaptively selected among mammals with distinct life history traits, like narrow habitat and smaller social groups, than that of gyrencephalic species(50). Distinct neurogenic potential of gyrencephalic and lissencephalic species has been attributed to the observed difference(50). Increased proliferative potential of basal progentior cells is necessary and sufficient to explain the gyrencephalic brains(50). The loss of such proliferative potential, which was likely an ancestral trait, might have caused inefficient neurogenic program in lissencephalic species. Our observation that the genes that had lost the synteny to CNEs in rat were involved in neuronal differentiation and were downregulated in fetal rat brain is largely coherent with the above proposal.

Altogether, our observations highlighted the link between genome order and the evolutionary dynamics of temporal gene expression pattern associated with mammalian brain development. The study also suggested that the genomic rearrangements, without any change in the genomic content, might impact the developmental trajectories and shape the evolution of phenotypes.

## Acknowledgement

Financial support to KSS from Department of Science and Technology (EMR/2015/001681) is duly acknowledged. MB and ML were financially supported by IISER-Mohali. KRC, YJ and HS thank ICMR, CSIR and SERB-DST agencies respectively for their fellowships.

## Methods

### Compilation of chromosomal position data

Human (hg19), rat (rn5), dog (camFam3), horse (equCab2) and cow (bosTau6) genome assemblies were used in the analysis. Conserved Non-coding Elements (CNEs) were taken from Marcovitz et al (Marcovitz et al.2016), which in turn were obtained by curating mammalian CNEs anchored to the human genome (hg19). Our choice of the aforementioned species and the CNE dataset was constrained by following considerations: i) We wanted sufficient evolutionary depth in the analysis and Marcovitz el al had considered 20 sequenced mammalian genomes to identify CNEs; 2) Since our analysis considered the chromosomal positions of CNEs and the genes, we only considered the genomes for which complete chromosome assemblies were available. For example, chromosome assemblies for the orders Cetacean, Chiroptera and Proboscidea etc. are not presently available; 3) In order to obtain the sufficient number of orthologous genes across species, we restricted our analysis to fewer mammalian lineages only. Considering multiple species would have compromised the total number of orthologous genes to start with.

We obtained the orthologue positions of human CNEs in query species using standard approach of mapping through LiftOver (https://genome-store.ucsc.edu/) chains at 0.95 mapping coverage(21). Finally, we compiled 114219 CNEs that were having orthologous positions in all 5 species. We independently obtained the table of orthologous genes across 5 mammals from Ensembl. Using CNE and gene tables, the list of nearest genes that were within 1Mb to the CNEs was obtained for human. The position of orthologous CNEs and genes in other mammals were assessed and CNE-gene pairs were classified as syntenic if the distance between the two was less than 1Mb in all 5 species and as non-syntenic if the CNE and the gene were >2Mb apart or were on different chromosomes in one of the species and remained within 1Mb in rest of the species. If there were multiple orthologues for the same gene, we took the nearest gene to the CNE on the same chromosome to ensure that a syntenic pair should not have classified as non-syntenic due to orthologue redundancy. The distance cut-off of 1Mb was determined based on distribution of number of CNE-gene pairs at different distance cut-offs. At around 1Mb, the overall distribution approached a plateau and the numbers did not increase significantly after that (Figure 1B). The lack of synteny cut-off of 2Mb ensured that CNE and the gene were distant at-least by 2 fold in their non-syntenic form when compared to their syntenic form. Larger distance cut-off was also likely to be robust against the annotation artefacts of gene coordinates. A flow-chart illustrating the overall strategy is given in Figure S1. All the data are available in supplementary data file.

To assess the genome assembly artefacts, we mapped the non-syntenic CNE-pairs to known problematic regions of rat genome (https://github.com/shwetaramdas/rataccessibleregions/). Out of 2711 CNE, only 3 CNEs (0.1%) and out of total 245 genes, only three genes (ABCC6, FOS, BNIP2; 1.2%) mapped to these regions. Exclusion of these regions was unlikey to change our claims. We further mapped the non-syntenic CNE-gene pairs of rn5 rat assembly to rn6 assembly. Out of 2711 CNE-gene pairs, 2667 pairs (98.4%) were successfully lifted over to rn6. Total 2227 (83.5%) pairs maintained non-sytenic status in rn6 too (Figure S7A). Removing the ambiguous pairs did not alter the signifcance of brain association (Figure SB). We also replaced the ENSEMBL orthologue information by *other_refseq* data in the above analysis to assess the correctnesss of orthologue mapping. The concordance of 83.5% and the persistence of brain association, therefore, confirmed that the observations presented in the article were robust against the technical artefacts of genome assembly and gene orthology.

### Analysis of genomic attributes

Chromosomal coordinates of repeat elements were downloaded from UCSC table browser. Repeat elements were mapped +/− 50kb around syntenic and non-syntenic CNEs and average value of enrichment in 2kb bins were plotted. For conservation analysis PhyloP scores of placental mammals (http://ccg.vital-it.ch/mga/hg19/phylop/phylop.html) were mapped +/− 1kb to CNE.

### Functional enrichment analysis

Gene Ontology and Mammalian Phenotype Ontology analysis was performed using GREAT (http://bejerano.stanford.edu/great/public/html/). Tissue specificity analysis was done using TSEA (http://genetics.wustl.edu/jdlab/tsea/), CSEA (http://genetics.wustl.edu/jdlab/csea-tool-2/). The tissue specificity index (pSI) score of a gene *i* in a tissue *k*, over the given tissues *j=1,2,…m* was calculated as per Dougherty et al (53) using following equation:

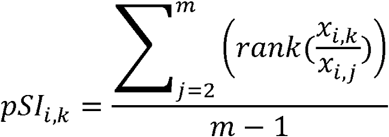

Where *x_i,1_* is the expression level of gene *i* in tissue 1 and *x_i,j_* is the expression level of gene i in tissue j. A stringent pSI cut-off of 0.05 was taken for the analysis. For syntenic gene set, we randomly sampled 245 genes (the size of non-syntenic gene set) from 4241 syntenic genes 100 times and plotted the mean and the standard error of the significance (-log10 of corrected p-value) of overlap between the candidate gene-sets and the tissues specific genes in the genome. The random sample of syntenic genes that exhibited most significant overlap with the brain-specific genes was taken for the expression specificity analysis among brain tissues across developmental stages.

Normalized gene expression data for developing cerebral cortex and heart of human, rat and sheep were taken from BRAINSPAN (human cortex; http://www.brainspan.org/static/download.html), GSE71148(human heart), Stead et al (rat cortex), GSE53512 (rat heart), Clark et al (sheep cortex) and GSE66725 (sheep heart). Average gene expression with 90% confidence intervals were plotted across development time course.

### Enhancer analysis

Regulatory potential of CNEs was assessed by mapping ChromHMM data obtained from Epigenome roadmap (http://egg2.wustl.edu/roadmap/web_portal/imputed.html#chr_imp) and ENCODE (http://hgdownload.soe.ucsc.edu/goldenPath/hg19/encodeDCC/wgEncodeBroadHmm/) projects onto CNEs. Enhancer coordinates from FANTOM (http://enhancer.binf.ku.dk/presets/) were also mapped to CNEs. Cumulative overlap across aforementioned three resources was calculated. Datasets for H3K4me1 methylation for fetal and post-natal/adult human tissues were obtained from Epigenome Roadmap (http://www.roadmapepigenomics.org/data/) with following accession IDs and age groups: fetal brain (E081, E082; 17GW), adult brain (E067, E068, E069, E071, E072, E073, E074; pooled 73Yr/75Yr/81Yr), fetal muscle (E089, E090; 15GW), post-natal muscle (E107; pooled 54Yr/72Yr) and fetal thymus (E093; 15GW), post-natal thymus (E112; 3Yr), fetal heart (E083, 91 days), post-natal heart (E95, E104, E105, pooled 3Yr/34Yr), fetal small intestine(E085, 15GW) and post-natal small intestine(E109, pooled 3Yr/30Yr). Fold-change over input DNA was used for aggregation plots. WashU epigenome browser was used for visualization. Motif analysis was performed through RSAT’s ‘peak-motif’ package (http://rsat.sb-roscoff.fr/peak-motifs_form.cgi) using JASPAR core matrices for vertebrate genomes. Syntenic CNEs were taken as background control sequences.

### Mapping of proxy GWAS SNPs

Total 251835 GWAS SNPs were obtained from GWASdb (http://jjwanglab.org/gwasdb). From this data, 71990 brain related SNPs were obtained by analyzing the HPO terms associated with brain associated phenotypes. We extended this data to total 533388 nearby SNPs (proxy) that were in linkage disequilibrium to 71990 brain related GWAS SNPs based on 1000 genome data using SNAP algorithm (https://personal.broadinstitute.org/plin/snap/index.php). Random null was prepared by picking CNE samples, of same sample size and CNE-gene distance as of non-syntenic set, from the syntenic set 1000 times and mapping SNPs to each of these samples. Number of CNEs with at-least one SNP was counted for each sample. The distribution of these numbers was regarded as random null. P-value was calculated as following:

Where B = number of re-sampling iterations (1000)
*k_b_* = Number of sampled syntenic CNEs having at-least one SNP.
*k* = Number of observed non-syntenic CNE having atleast one SNP.

### Virtual 4C data

SRA files of HiC datasets for fetal and adult brains were obtained from GSE77565 and GSE87112. Datasets were processed using HiCUP (https://github.com/theaidenlab/juicer/wiki/HiCCUPS) and contact maps were normalized using iterative correction and eigen vector decomposition (ICE) method (https://github.com/hiclib/iced). TSS in each CNE-gene pair was taken as bait (reference point) and its intra-chromosomal interactions were obtained from HiC matrices. Loess regression line was fit to the HiC counts as a function of genomic distance from the bait. Signifcant interactions with the bait were identified by applying cut-off of 3-standard deviation distance from the regression line (54).

### DNA breakpoint analysis

We obtained 552 germline breakpoints associated with congenital disorders having brain abnormalities and 68018 somatic cancer breakpoints from van Heesch et al (55). The matching RNA-seq data of perpheral blood of patient and the mother were obtained from European Nucleotide Archive(https://www.ebi.ac.uk/ena) with the accession IDs ERX358048 and ERX358046 respectively. Total 2061 evolutionary DNA break-points for rodents were taken from Bourque etal (2004 & 2006), Larkin etal and Lemaitre et al (56-59). These breakpoints were then mapped onto inter-spacer regions between CNE and the nearest gene-TSS. The random null was obtained by picking CNE-gene pairs, of same sample size and CNE-gene distance as of non-syntenic set, from the syntenic set 1000 times and mapping the breakpoint in the inter-spacer regions. Number of CNE-gene pairs with at-least one break-point in between was counted for each sample. The distribution of these numbers was regarded as random null. P-values was calculated using equation as in GWAS SNP analysis. The break-point distances from the ancestor were obtained from Luo et al(25).

### Mammalian traits

Status of morphological traits in 5 mammals were obtained from project ID P773 of Morphobank database (https://morphobank.org/). Traits that exhibited same status atleast in 3 of the mammals including rat, but showed a different status in human were classified as independently modified traits in human. Similarly the traits that had same status in atleast 3 species including human, but changed status in rat were denoted as independently modified traits in rat.

### Availability of data

All datasets presented in this article are available as supplementary data.

## Supplementary Figure Legends

**Figure S1.** Flowchart illustrating overall strategy to obtain syntenic and non-syntenic CNE-gene pairs.

**Figure S2.** Enrichment of (a) Mammalian Phenotype Ontology (MPO) terms, and (b) Bgee anatomical terms in syntenic (left panel) and non-syntenic (right panel) gene-sets. P-values were corrected using Benjamini-Hochberg method.

**Figure S3.** Extension of Epigenomic analyses of syntenic and non-syntenic CNEs. (A-B) H3K4me1 ChIP enrichment (over inut DNA) around syntenic (grey) and non-syntenic (red) CNEs in (a) fetal, and (b) post-natal tissues. (C) Strategy used to assess the CNE-TSS interactions using HiC datasets of human fetal and adult brains. All-to-all HiC interactions were filtered for TSS-to-all interactions for the genes in syntenic and nonsynteic sets. The resultant data was analogous to 4C and was analyzed using method atypical for 4C analysis. Loess regression line was fit to 4C counts as a function of genomic distance from the refrence point (TSS in this case). The distance of 3-standard deviation from this regression line was taken as signicance cut-off for the interactions impinging onto TSS. (D) Example of CNE-TSS interactions identified in the non-syntenic set. Upper and lower panels represent fetal and adult brain data. Red line: 4C signal; grey line: H3K4me1; Black line: Loess fit; Dotted line: 3-standard deviation cut-off for signicance.

**Figure S4.** Sequence motif enrichment analysis for non-synteic CNEs. ‘Peak-motifs’ from RSAT was used to identify over-represented sequence motifs in the non-synteic CNEs while taking syntenic CNEs as background control. (a) Sequence motifs, their e-values and the matching transcription factors (TFs) from JASPAR. (b) Tissue specificity analyses of TFs. Red bars represent the tissues, wherein the TFs exhibit significant specificity (P<0.05). (c) Time course gene expression of TFs during human brain development. Red curve represents average expression of TFs and grey colour denotes 90% confidence interval.

**Figure S5.** Independently modified traits in rat and human. (A) bar plot representing rat-to-human ratio for the proportion of traits independently modified in each trait class. (B) List of traits that were independetly modified in rat.

**Figure S6.** Analysis of gain of synteny instances in human. Gain of synteny instances were identified using the strategy shown. No enrichment was found in the GO process terms. The most enriched GO term is shown.

**Figure S7.** Comparison of non-syntenic (rat) CNE-gene pairs in rn5 and rn6 genome assemblies. (A) Shown is the scatter plot of CNE-gene distances (log10 scale) of the non-syntenic set in rn5 and rn6 assemblies. (B) Gene Ontology enrichment analysis of genes that had lost synteny consistently in both assemblies (83.5% of total). P-values are adjusted using Benjamini-Hochberg method.

